# Viral pIC-pocketing: RSV sequestration of eIF4F Initiation Complexes into bi-phasic biomolecular condensates

**DOI:** 10.1101/2023.04.14.536931

**Authors:** Fatoumatta Jobe, James T. Kelly, Jennifer Simpson, Joanna Wells, Stuart D Armstrong, Matt Spick, Emily Lacey, Leanne Logan, Nophar Geifman, Philippa Hawes, Dalan Bailey

## Abstract

Orthopneumoviruses characteristically form membrane-less cytoplasmic inclusion bodies (IBs) wherein RNA replication and transcription occur. Herein, we report a strategy whereby the orthopneumoviruses sequester various components of the eiF4F <u>I</u>nitiation <u>C</u>omplex machinery into viral IBs to facilitate translation of their own mRNAs; p<u>IC</u>-pocketing. Mass spectrometry analysis of sub-cellular fractions from RSV-infected cells identified significant modification of the cellular translation machinery; however; interestingly, ribopuromycylation assays showed no changes to global levels of translation. Electron micrographs of RSV-infected cells revealed bi-phasic organisation of IBs; specifically, spherical “droplets” nested within the larger inclusion. Using correlative light and electron microscopy (CLEM), combined with fluorescence in situ hybridisation (FISH), we showed that the observed bi-phasic morphology represents functional compartmentalisation of the IB and that these domains are synonymous with the previously reported inclusion body associated granules (IBAGs). Detailed analysis demonstrated that IBAGs concentrate nascent viral mRNA, the viral M2-1 protein as well as many components of the eIF4F complex, involved in translation initiation. Interestingly, although ribopuromycylation-based imaging indicates the majority of viral mRNA translation likely occurs in the cytoplasm, there was some evidence for intra-IBAG translation, consistent with the likely presence of ribosomes in a subset of IBAGs imaged by electron microscopy. The mechanistic basis for this pathway was subsequently determined; the viral M2-1 protein interacting with eukaryotic translation initiation factor 4G (eIF4G) to facilitate its transport between the cytoplasm and the separate phases of the viral IB. In summary, our data shows that IBs function to spatially regulate early steps in viral translation within a highly selective biphasic liquid organelle.

## Introduction

Orthopneumoviruses, including human and bovine respiratory syncytial virus (RSV) as well as pneumonia virus of mice (PVM), are taxonomically classified within the *Pneumoviridae* family and *Mononegavirales* order of negative-strand RNA viruses [1, 2]. Human (h) and bovine (b) RSV are globally prevalent and associated with respiratory tract infections; posing significant health and economic burdens [3, 4]. Despite their host specificity, the two viruses are genetically and antigenically similar and share orthologous mechanisms of replication and pathogenesis [5-7].

Their small 15 kb non-segmented single stranded RNA genomes encode 11 proteins from 10 transcriptional units [8], existing as a helical ribonucleoprotein (RNP) complex, together with multiple copies of the nucleoprotein (N) [9]. Encoded proteins include those required for autonomous genome replication, mRNA transcription [10] and host immune regulation [11-13]. Due to their negative sense genome, the incoming RNP complex is associated with the RNA-dependent RNA polymerase (L), the phosphoprotein (P) and viral transcription factor (M2-1) [14, 15] required for replication and transcription. Like many negative sense RNA viruses, genome replication and transcription occur in cytoplasmic membrane-less replication complexes, called inclusion bodies (IBs) [16-18], with the RSV N and P proteins being key drivers of the liquid-liquid phase separation (LLPS) needed for IB formation [16, 19].

IBs formed during infection have dynamic properties and host a multitude of viral-viral and viral-host protein interactions underpinning a diverse range of functions [20-23]. The best characterised is the concentration of viral proteins for the spatio-temporal compartmentalisation of viral RNA replication and mRNA transcription [16-18, 24]. Furthermore, a number of host cell proteins involved in the antiviral response (NF-κB subunit p65 [18], MAVS and MDA5 [25], phosphorylated p38 and O-linked *N*- acetylglucosamine transferase [26]) are sequestered into IBs to prevent their function. Protein phosphatase 1 (PP1) recruitment to the IB has also been shown to be involved in important post-translational regulation steps in the viral life cycle [22].

We and others have also previously shown that IBs are highly organised, with newly synthesised mRNA and the RNA-binding M2-1 protein [18, 27] further compartmentalised into sub-IB droplets, termed inclusion body-associated granules (IBAGs) [17, 18, 27]. The presence of poly(A)-binding protein (PABP) and the translation initiation factor eIF4G in these mRNA-rich droplets [17] suggests a role in translation initiation [28]. Although it has been suggested that IBAG contents, are released into the cytoplasm for viral translation to occur [17], there is little information available on the ultimate site for translation of RSV viral mRNAs.

To this end we used mass spectrometry to examine the spatio-temporal re-organisation of host and viral proteins during RSV infection and identified significant regulation of proteins involved in RNA metabolism and translation. Subsequent ribopuromycylation assays combined with immunofluorescence microscopy identified sites of active translation and sub-cellular localisation of the translation initiation machinery in infected cells. These changes were correlated to viral modification of the intracellular environment using electron microscopy and correlative imaging. In a limited number of IBAGs ribosomes were present, consistent with evidence for intra-IBAG translation. We also observed that components of the eIF4F complex both colocalised and co-immunoprecipitated with RSV M2-1 and viral mRNA in IBAGs, which appear as more soluble IB microdomains within IBs. In summary, our findings represent a novel strategy, pIC-pocketing, by which orthopneumoviruses can hijack components of the cellular cap-dependent translation machinery into viral IBs and broadly contributes to our understanding of RSV replication.

## Results

### The RSV proteome is predominantly membrane associated, with infection resulting in modification of RNA metabolism and translational machinery in the cell

To examine proteomic changes to the intracellular microenvironment following RSV infection we performed sub-cellular fractionation of uninfected and hRSV infected cells at 24 and 48 hours post infection (hpi). Initial experiments identified satisfactory fractionation of these compartments, validated by western blot (Supplemental Fig. 1A). Cytosolic and membrane fractions were then harvested from biological replicates (n=5) and analysed by label free quantitative mass spectrometry (Supplemental Fig. 1B). Principal component analysis (PCA) of the thirty samples showed that component one (capturing the largest part of the variation, plotted on the x-axis) showed differentiation between the cytosol and membrane, with this continuing over the duration of the experiment (Fig. 1A). The second component of the PCA captured variation corresponding to time/infection (plotted on the y-axis). In the majority, comparison of the ion abundancies for the successfully detected proteins from the RSV proteome (NS1, N, P, M, G, F, M2-1 and L) demonstrated that RSV infection is predominantly associated with the membrane fraction, the only exception being NS1 and L at 24 hpi (Fig. 1B). Interestingly, the most abundant viral proteins in the membrane fraction at 48 hpi were N, P and M2-1, despite the lack of well recognised transmembrane domains in these proteins. Comparison of all detected proteins within the membrane and cytosolic fractions highlighted the significant and robust detection of the virome in the membrane fraction, as well as significant differential regulation of hundreds of cellular proteins (Fig. 2C and 2D). Pathway analysis of all proteins with an adjusted FDR p-value of <0.01 highlighted significant enrichment of proteins involved in RNA metabolism and infectious disease within the membrane fraction (Fig. 1E) and multiple overlapping translation pathways within the cytosol fraction (Fig. 1F). For Reactome pathways ‘RNA metabolism’ (R-HSA:8953854) and ‘Translation’ (R-HSA:72766) (highlighted in green in Fig. 1E and 1F) we demonstrated that the majority of the positively associated proteins were present with the negative fold change (Fc) component of the volcano plot (Fig. 1G and 1H respectively), indicating a global trend for downregulation of these proteins. When we repeated fractionations and examined a number of proteins well established to play a role in translation by Western blot (Fig.1I and 1J), including three identified in our mass spectrometry pathway analysis (Fig.1I; eIF3A, RPL3 and RPS6; R-HSA:72766) we did not observe gross downregulation of these proteins in the whole cell lysate, consistent with the lower fold changes observed (median Fc of -0.89 log2; n=54 detected proteins in R-HSA:72766) (Fig. 1I). However, we did see slightly modified expression of eIF4G, eIF3A, RPS6 and RPL3 when comparing the membrane and cytosol fractions (Fig. 1I and 1J). To help elucidate the nature of these modifications we turned our attention to immunofluorescence based sub-cellular localisation studies.

**Figure 1.**
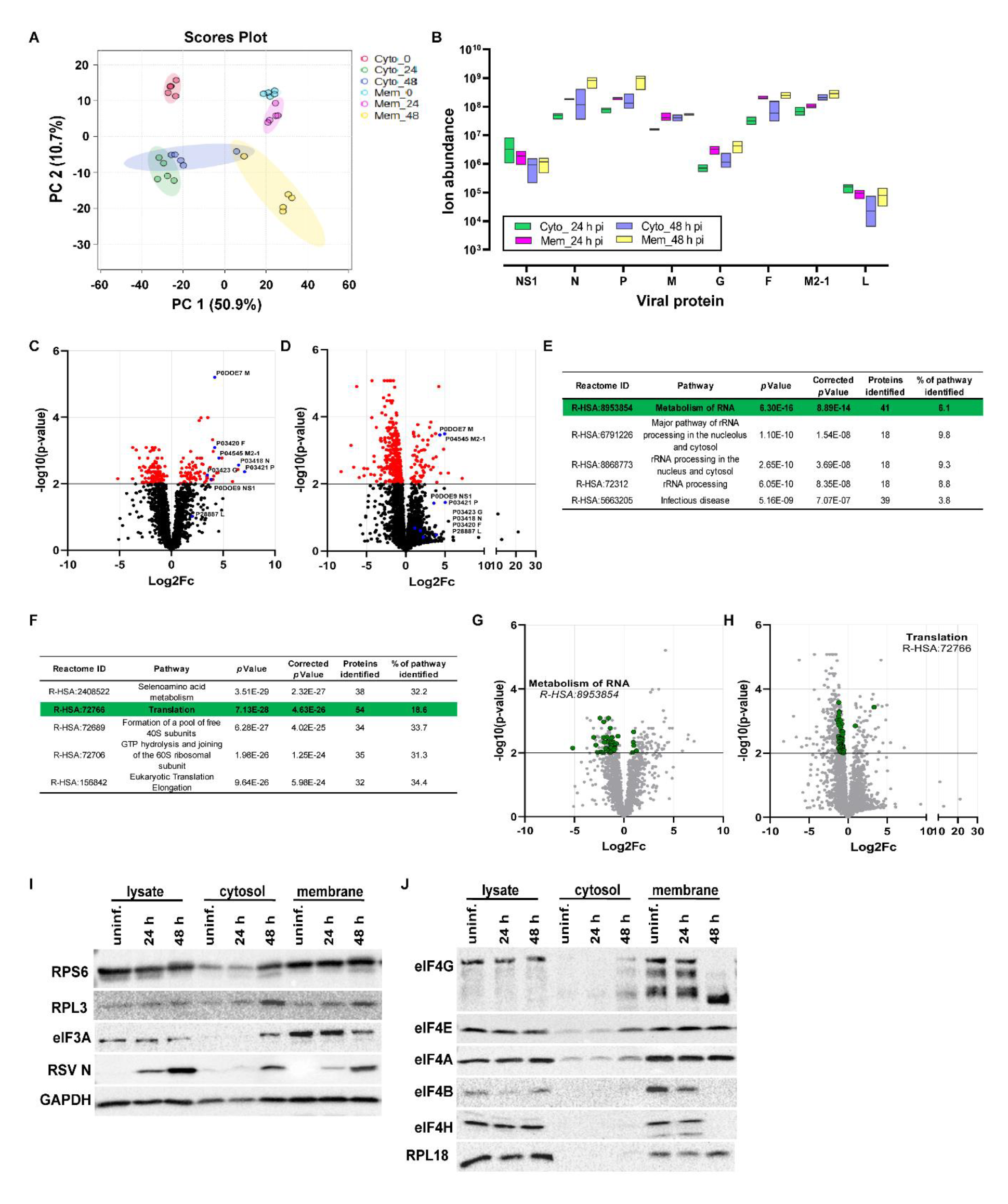
The RSV proteome is predominantly membrane associated, with infection resulting in modification of RNA metabolism and translational machinery in the cell. (A) Quintuplicate (n=5) membrane (Mem) and cytoplasmic (Cyto) fractions from either mock or hRSV-infected (24 and 48 h p.i.) cells were isolated using the Thermofisher Mem-PER Plus Membrane protein Extraction Kit and analysed by label free quantitative mass spectrometry. Protein fractions were run on SDS -PAGE and subjected to in-gel tryptic digestion. Peptides were analysed on a Thermo Q-Exactive mass spectrometer. Normalised and log transformed ion abundancy data from these 30 fractions was analysed by principal component analysis (PCA) to show the distribution of the samples analysed when the feature space is reduced to two dimensions. The PCA axes show the first and second most important components in terms of the reduced space along which the samples show the largest variation. The units of the axes are percent variation in the overall dataset explained by the two components. (B) Ion abundancies for positively detected hRSV proteins are shown in the respective membrane (Mem) and cytoplasmic (Cyto) fractions at both 24 and 48 hpi. Each floating boxplot represents the minimum and maximum abundancies observed from the 5 biological replicates, with the line representing the mean. NS2, M2-2 and SH were not detected. (C and D) Volcano plots reflecting comparison of ion abundancies between mock and infected membrane (C) and cytosolic (D) fractions over time. Fold changes (log2) and p-values (-log10 of the false discovery rate [FDR]) are plotted, with proteins meeting the criteria for pathway analysis (FDR-adjusted p-value of <0.01) highlighted in red. The eight successfully detected hRSV proteins are separately highlighted in blue in both C and D together with their accession numbers and protein names. All other proteins are represented by black symbols. (E and F) Pathway analysis of detected proteins with an adjusted FDR p-value of <0.01 was conducted seperately for the membrane (E; n= 203 proteins) and cytosolic (F; n= 407 proteins) fractions against the Reactome pathway database (REACTOME_Pathways_25.05.2022). The top 5 pathways identified (Reactome IDs ranked by Terms p-value as well as Bonferroni step down adjusted p-value) are tabulated, together the number and total percentage of proteins from this pathway successfully identified. (G) The 41 successfully identified proteins from the ‘RNA metabolism’ pathway (R-HSA:8953854; highlighted in green in E) are shown in a volcano plot of differentially regulated proteins within the membrane fraction of infected cells from C. (H) The 54 successfully identified proteins from the ‘Translation’ pathway (R-HSA:72766; highlighted in green in F, are shown in a volcano plot of differentially regulated proteins within the cytoplasmic fraction of infected cells from D. In both G and H, all other proteins are greyed out. (I and J) Immunoblots show levels of the indicated proteins in whole cell lysates (lysate), in cytosolic (cytosol) or membrane fractions prepared from uninfected (uninf.) or from hRSV-infected (24 or 48 h p.i.) A549 cells.

**Figure 2.**
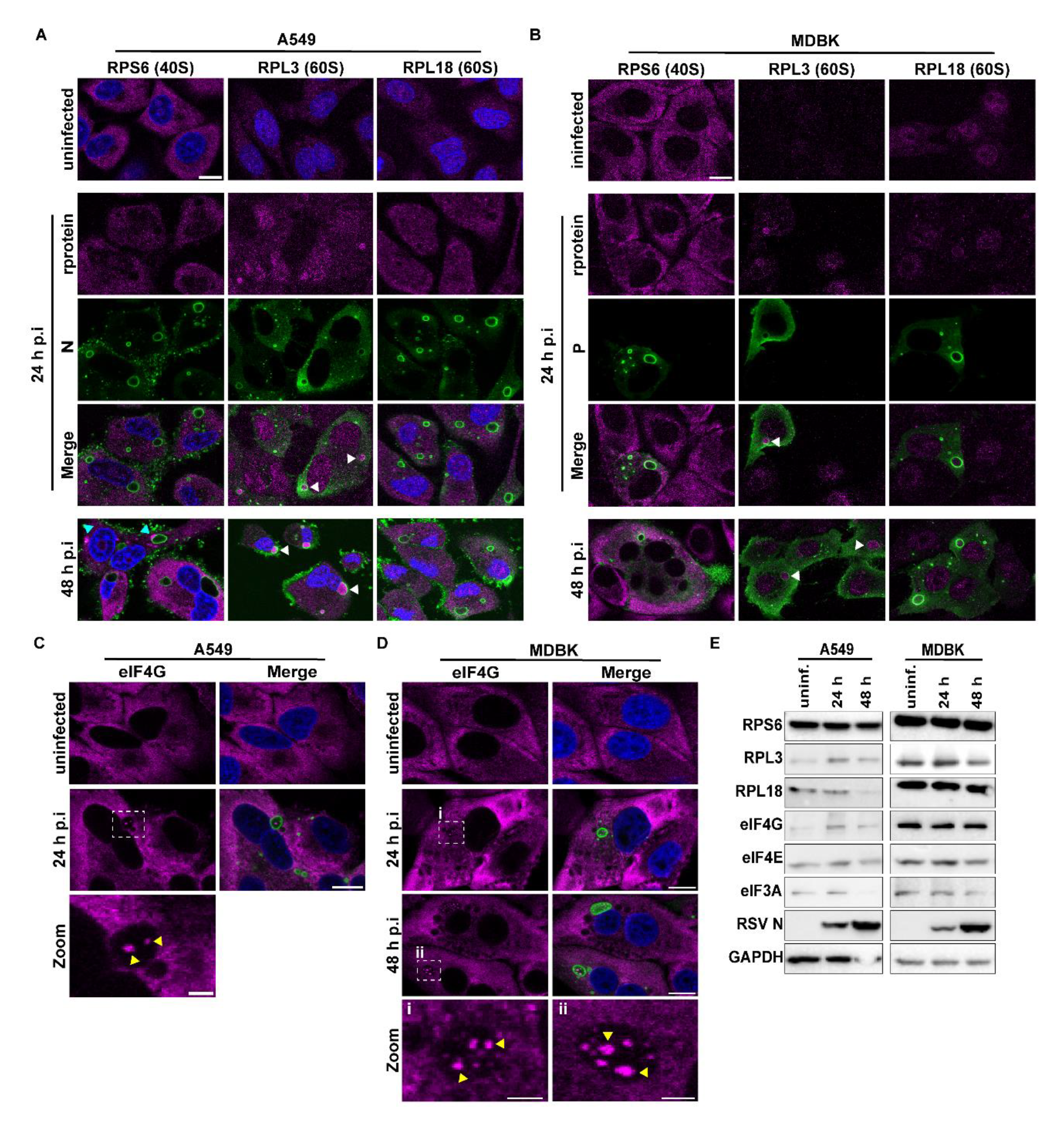
Spatio-temporal characterisation of ribosomal subunits and translation initiation scaffold protein, eIF4G. (A and B) Uninfected or cells infected with RSV (A; hRSV-infected A549 cells and B; bRSV-infected MDBK cells) at an MOI of 2 for 24 or 48 h were fixed, permeabilised and immuno-stained with antibodies against RSV N or P protein (green), ribosomal subunits, RPS6, RPL3 and RPL18 (magenta), and nuclei stained with DAPI (blue). Images are representative of data from three (A) and two (B) independent experiments and were obtained using a Leica Stellaris confocal microscope. Scale bar, 10 µm. (C) A549 and (D) MDBK cells respectively infected with hRSV or bRSV at an MOI of 1 for the indicated times, were fixed and immuno-stained for eIF4G (magenta) and RSV N (green) followed by confocal imaging. Zoom panels are enlarged images of the boxed areas; unlabelled in C and i and ii in D. Scale bar, 10 µm in main and 2 µm in zoom. (E) Immunoblots show levels of the indicated proteins in whole cell lysates prepared from uninfected (uninf.) or from hRSV-or bRSV-infected (24 or 48 h p.i.) A549 or MDBK cells, respectively.

### 60S Ribosomal protein subunit, RPL3, and eukaryotic translation initiation factor, eIF4G, are spatially reorganised into RSV IBs

Based on the results of our pathway analysis we sought to explore the spatio-temporal organisation of components of the cellular translation machinery during infection. Immunofluorescence (IF) microscopy was performed to examine the location of protein components of both 40S (RPS6) and 60S (RPL3 and RPL18) ribosomal subunits. IF analysis showed minimal changes to the subcellular distribution of RPS6 and RPL18 following infection of human (A549) cells (Fig. 2A), although both proteins were excluded from cytoplasmic viral IBs (ring structures of N staining; green). Interestingly, however, we observed recruitment of RPL3 into IBs in infected cells (Fig. 2A; white arrow head) similar to our previous observations for NF-κB subunit p65 [18]. By 48h p.i., we noted that the distributions of RPL3 and RPL18 remained unchanged, however punctate structures of RPS6 (cyan arrowhead), were sometimes found in close proximity to IBs. Consistent findings were also obtained in bRSV infected bovine (MDBK) cells (Fig. 2B) providing further evidence of orthologous replication strategies between these two orthopneumoviruses. Building on these findings and previous reports of eIF4G, total polyadenylated (PolyA) and viral mRNA association with RSV IBs [17], we next examined the location of the cellular translation initiation machinery during infection. We first confirmed the recruitment of eIF4G, a key scaffolding component of the eIF4F translation initiation complex, involved in the recruitment of mRNA to the ribosome. In uninfected A549 and MDBK cells, eIF4G was distributed throughout the cytoplasm (Fig. 2C and D). However, following infection, eIF4G was recruited into a sub-set of IBs and spatially concentrated in microdomains (yellow arrow heads) or IBAGs, as previously reported.

Despite the reorganisation of this machinery and our pathway analysis whole cell lysate analysis by Western blot showed comparable levels of these proteins (and others involved in translation; eIF4E and eIF3A) between infected and uninfected lysates, except for RPL18 at 48 h p.i when some cytopathic effects (cpe) were emerging (Fig. 2E), consistent with our analysis of cellular fractions (Supplemental Figure 1D). Together, these findings indicate the spatial dysregulation of selected ribosomal subunits and translation initiation machinery in orthopneumovirus infected cells.

### RSV IBs are biphasic and recruit components of the eIF4F complex into specialised micro-domains

To facilitate electron microscopy analysis, we next validated the use of bovine RSV infection of Vero cells as a suitable model for orthopneumovirus infection. Consistent with the results in Fig. 2, we observed selective recruitment of eIF4G into microdomains within IBs, with no reduction in the level of ribosomal subunits or eIF4F complex proteins detectable by Western blot analysis (Fig. 3A). Following this, we assessed the localisation of other eukaryotic translation initiation factors (eIF4A, 4A1, 4B and 4H) in these cells. Briefly, the eIF4F complex with eIF4G acting as the scaffold is essential in the initiation step of protein synthesis, recruiting mRNA to the 43S preinitiation complex through multiple interactions depicted in Fig.3B, ultimately leading to 80S ribosome formation [29]. In the absence of infection, the eIF4F complex proteins eIF4A, 4A1, 4B and 4H are distributed in the cytoplasm with varying degrees of nuclear staining (Fig. 3C). As for eIF4G, infection induced their recruitment to sub-IB microdomains (Fig. 3D), potentially indicating pre-assembly of the eIF4F complex within IBAGs.

**Figure 3.**
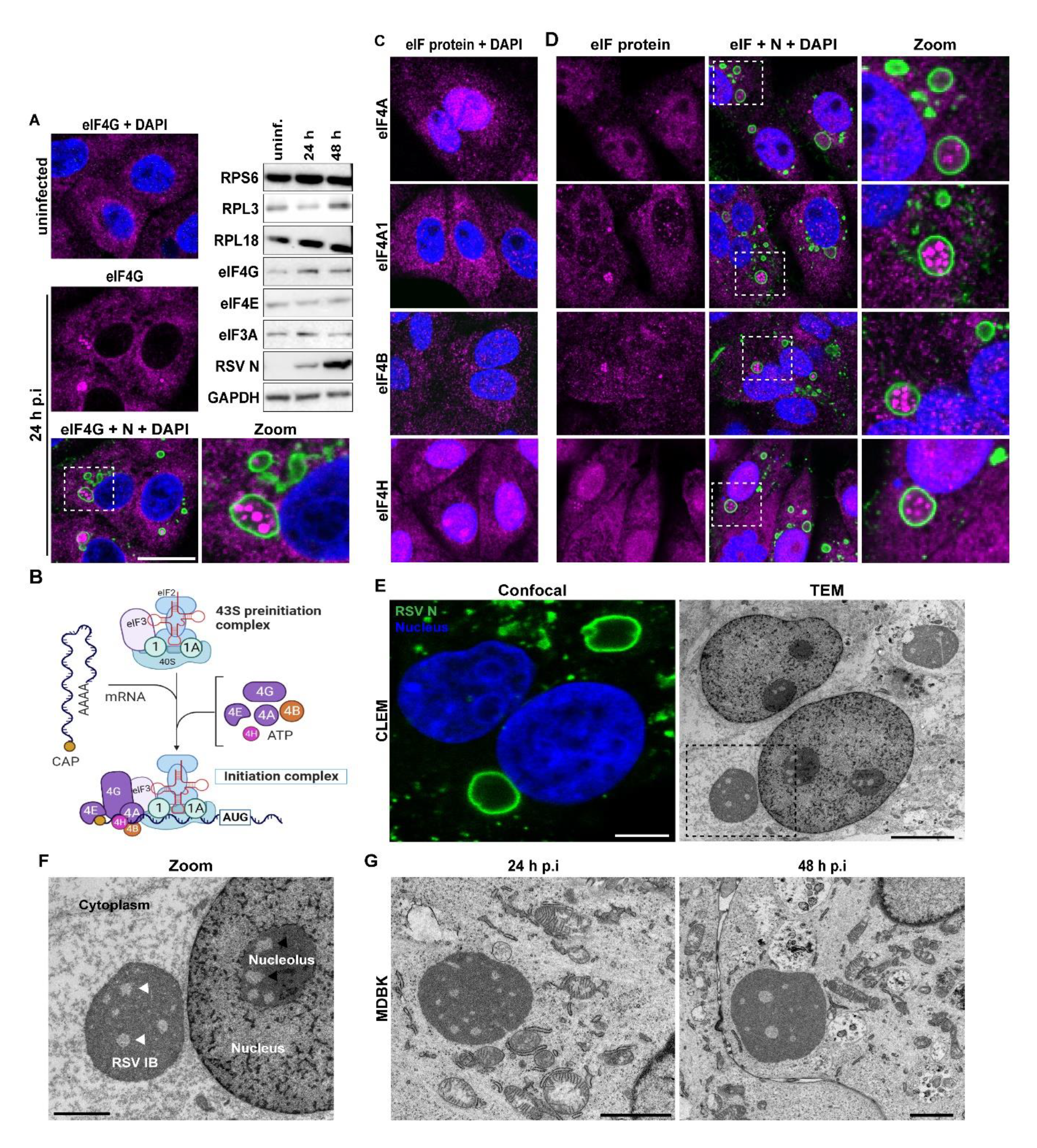
eIF4F components are recruited into microdomains within bi-phasic IBs. (A) Vero cells were left uninfected or infected with bRSV at an MOI of 1 for 24 h. Cells were then fixed and immuno-stained for eIF4G (magenta), RSV N (green) and nuclei stained with DAPI (blue) for confocal analysis. Scale bar 20 µm. Immunoblots show levels of the indicated proteins in Vero cell lysates – uninfected or infected with bRSV at an MOI of 1 for 24 or 48 h. (B) A schematic representation of the association of 43S preinitiation complex with 5’ capped mRNA and components of the eIF4F protein complex to form the initiation complex. Adapted from “Protein Translation Cascade” template in BioRender.com (C) Uninfected and (D) bRSV infected Vero cells (MOI of 1 for 24 h) were fixed and stained for the indicated eIF4F component (magenta), RSV N (green) and nuclei (blue). Enlarged images of the boxed areas in merged images of A and D (eIF4G + N + DAPI), are shown in the “zoom” panel. (E) Correlative light electron microscopy (CLEM). Vero cells infected with bRSV at an MOI of 1 for 24 h were fixed and immuno-stained with antibodies against RSV N (green) and nuclei stained with DAPI for confocal microscopy. Following confocal imaging, cells were prepared for transmission electron microscopy (TEM) and the same cells imaged by confocal (left) identified and visualised by TEM (right). Scale bars, 5 µm. (F) A higher magnification of the IB/area boxed in E. Arrow heads indicate areas of reduced electron density within the IB and nucleolus. Scale bar, 2 µm. (G) MDBK cells infected with bRSV for the indicated times, fixed in glutaraldehyde and prepared for TEM. Scale bars, 2 µm.

We previously reported that bovine RSV IBs are membrane-less liquid organelles that form by liquid-liquid phase separation (LLPS) [18]. Correlative analysis by CLEM again highlighted the ultrastructure of the IBs (Fig. 3E) and the significant modification of the cytoplasmic environment, when compared to uninfected cells (Supplementary Fig. 2B). Interestingly, however, our detailed analysis of TEM images showed that a sub-set of RSV IBs are bi-phasic, with multiple areas of reduced electron density found within the IB itself (Fig. 3E and 3F). We hypothesised that these are synonymous with the IBAG microdomains observed by IF in Fig. 3A and D. This biphasic nature to the IB, to our knowledge unique in composition and structure within the characterised liquid organelles of non-segmented RNA viruses; shows remarkable similarity with the spatial organisation of the nucleolus, a characteristic cellular condensate (Fig. 3F). Of note, microdomains of reduced electron density within the nucleolus are fibrillar centres where ribosomal RNA transcription and pre-ribosomal RNA processing occur [30]. It has been suggested that the morphology of biphasic condensates is primarily governed by sequence-encoded and composition-dependent protein RNA interactions [31] and likely holds true for our spatially organised RSV IBs. Examination of bRSV IBs formed in MDBK cells (uninfected cells in supplementary Fig. 2C) also showed similar organisation further validating the suitability of Vero cells as models to study orthopneumovirus biology.

### Biphasic morphology of RSV IBs correlates with functional stratification of the condensate

RSV infection characteristically induces modification of the cellular micro-environment, with viral RNA replication and mRNA transcription taking place inside IBs. However, there is little evidence available on the spatial organisation of RSV mRNA translation. Previously, we showed that nascent viral RNA and RSV M2-1 protein co-localised in IBAGs during infection [18]. Here, we first examined the potential colocalization of M2- 1 with eIF4G and 4A1 and found strong colocalization of M2-1 with both proteins within IBs, although M2-1 was also found throughout the IB (Fig. 4A). This data confirms that M2-1 rich IBAGs are synonymous with the sub-IB micro-domains that concentrate eIF4F components. Next, we sought to confirm the identity of the sub-IB microdomains identified by TEM through correlation with IF staining of IBAG markers. To identify the best marker of IBAGs for CLEM, we combined fluorescence *in situ* hybridisation (FISH) staining with IF microscopy. We observed concentration of NS1 and N mRNA in small foci (white arrow heads) inside a sub-set of IBs, in a pattern consistent with IBAGs (Fig. 4B). PolyA FISH and M2-1 co-stain confirmed mRNA accumulation in IBAGs (white arrows in Fig. 4C) and confirmed polyA FISH as the best marker of IBAGs, likely due to its relative higher abundance, when compared to specific viral mRNAs and/or proteins. To develop a deeper understanding of IB morphology 3D reconstruction analysis was subsequently used to quantify IB and IBAG volumes. M2-1 (Fig. 4C) or P (supplementary Fig. 2D) signals were used as source for IB identification and PolyA for IBAGs (Figure 4C; insets). These analyses demonstrated the heterogeneity in size and number of IBAGs (Fig 4D; images i-v and left graph; also supplementary Fig. 2D). In addition, it provided evidence that IB size is not a determinant of IBAG number and volume. For example, IB i, 24.77 µm³ in volume, has more numerous and larger IBAGs than the larger IB v (50.48 µm³), likely consistent with previous findings that these structures are highly transient in nature [17]. Nevertheless, IBAGs were mostly present in IBs with diameter above 4 µm (Fig 4D; right graph). Having established polyA FISH as the best marker for IBAG detection, we used this approach for CLEM analysis of infected cells. Importantly, these experiments confirmed that fluorescent polyA signals (IBAGs) did indeed correlate with sub-IB microdomains of reduced electron density (white arrows) in both bRSV-infected Vero (Fig. 4E) and hRSV-infected A549 cells (Fig. 4F), validating our earlier hypothesis and confirming the bi-phasic nature of IBs relates to RNA-and RNA-binding protein-rich microdomains.

**Figure 4.**
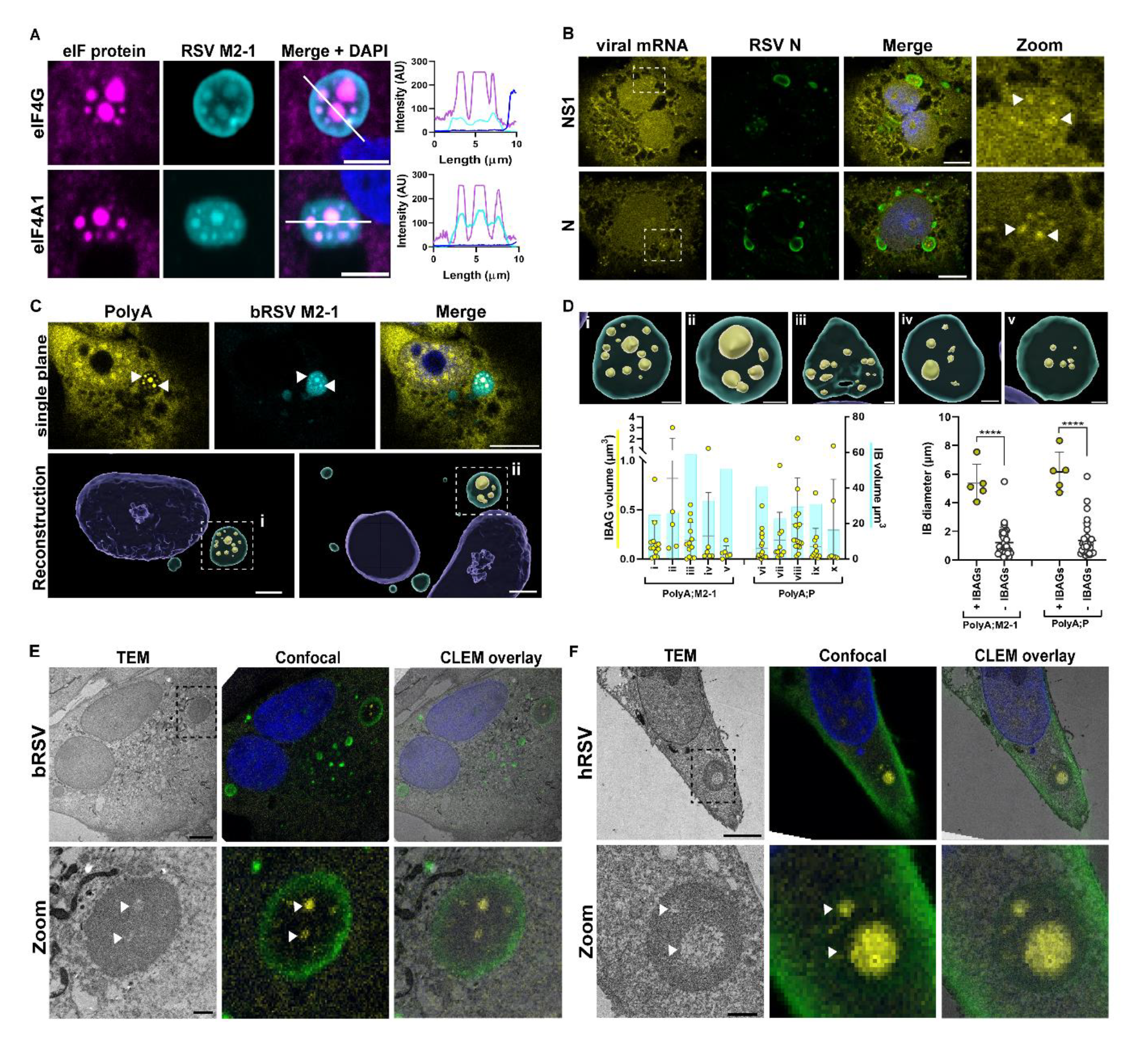
M2-1 and viral mRNA concentrate in the differentially phase separated microdomains. Vero cells infected with bRSV at an MOI of 1 were fixed at 24 h p.i. (A) Cells were immuno-stained for eIF4G (magenta), RSV M2-1 (cyan) and nuclei stained with DAPI (blue) for confocal image analysis. Scale bar, 5 µm. Graphs show fluorescent intensity profiles along the white lines drawn in the “merge + DAPI” panel. (B and C) FISH analyses were performed (as described in the methods) with specific biotinylated probes to detect the indicated viral mRNAs (B; yellow) or total polyadenylated (PolyA) mRNA (C; yellow). Negative controls included uninfected cells stained with the same probes or bRSV infected cells stained with probes against VSV G mRNA. Co-immunostaining was performed with RSV N (green) in B and M2-1 (cyan) in C. Presented images are single-plane confocal images taken in the median section of the cells. Scale bars, 10 µm. Arrow heads indicate sites of mRNA and M2- 1 concentration. Bottom panel of C – Imaris reconstruction of confocal image Z-stacks of the same cell from the top panel (left) and another example (right). Scale bars, 3 µm. (D) Enlarged images of the IBs boxed in C (i and ii) are shown with other examples (iii to v). Scale bars, 1 µm. Left graph – volume of the IBs in the panel above (represented as cyan bars) and volumes of individual IBAGs identified within them (presented as yellow dots), also quantified using the Imaris software, as described in the methods. IBs i-v were stained for PolyA and M2-1, and vi-x with PolyA and P. Right graph – Longest diameter of IBs containing identifiable IBAGs (+ IBAGs) and those without (- IBAGs), also quantified using the Imaris software. An unpaired *t*-test was used for statistical analysis; ****, P<0.0001. (E and F) CLEM analysis of (E) Vero cells infected with bRSV and (F) A549 cells infected with hRSV. TEM was done of the same cells following confocal imaging for PolyA mRNA (yellow) and RSV P (green) and images merged in the CLEM overlay panels. Scale bars, 5 µm. Zoom panels are higher magnification images of the IBs boxed in the panels above. Scale bars, 1 µm.

### Biphasic IBAGs are functional sites containing ribosomes and ribopuromycylated products

Detailed analysis of TEM images of bi-phasic IBAGs indicated the putative presence of ribosomes in a sub-set of IBAGs in both bRSV-infected Vero and MDBK cells (black arrows and i in Fig. 5A). Importantly, these were not always present, with many IBAGS having little evidence for ribosomal colocalization (Fig. 5A; white arrows and ii). Nevertheless, these observations, in combination with our earlier findings on eIF4F, viral mRNA and RPL3, suggest that viral translation might be occurring in select IBAGs. To assess this, we used ribopuromycylation to interrogate sites of translation during infection. Co-immunostaining for nascent polypeptide-incorporated puromycin (PRM) and eIF4G (Fig. 5B), polyA (Fig. 5C) or RSV P (Fig. 5D) in bRSV-infected cells revealed the presence of active translation in some IBAGs, although the majority of the labelled signal was evenly distributed within the cytoplasm (Fig. 5B-D). Unsurprisingly, varying PRM signals were observed within IBs, from significant (Fig. 5C, e.g.1) to none (Fig. 5C, e.g.2), correlating with our TEM observations. A broader ‘population level’ analysis by coomassie staining and immunoblotting of whole cell lysates revealed that, unlike many RNA virus infections, global levels of protein synthesis remained, intriguingly, unaffected by RSV infection. Together, these findings demonstrate that IBAGs, areas of reduced electron density in EM, frequently contain ribosomes and can act as functional sites for translation, albeit without wholesale effects on translation within the infected cell.

**Figure 5.**
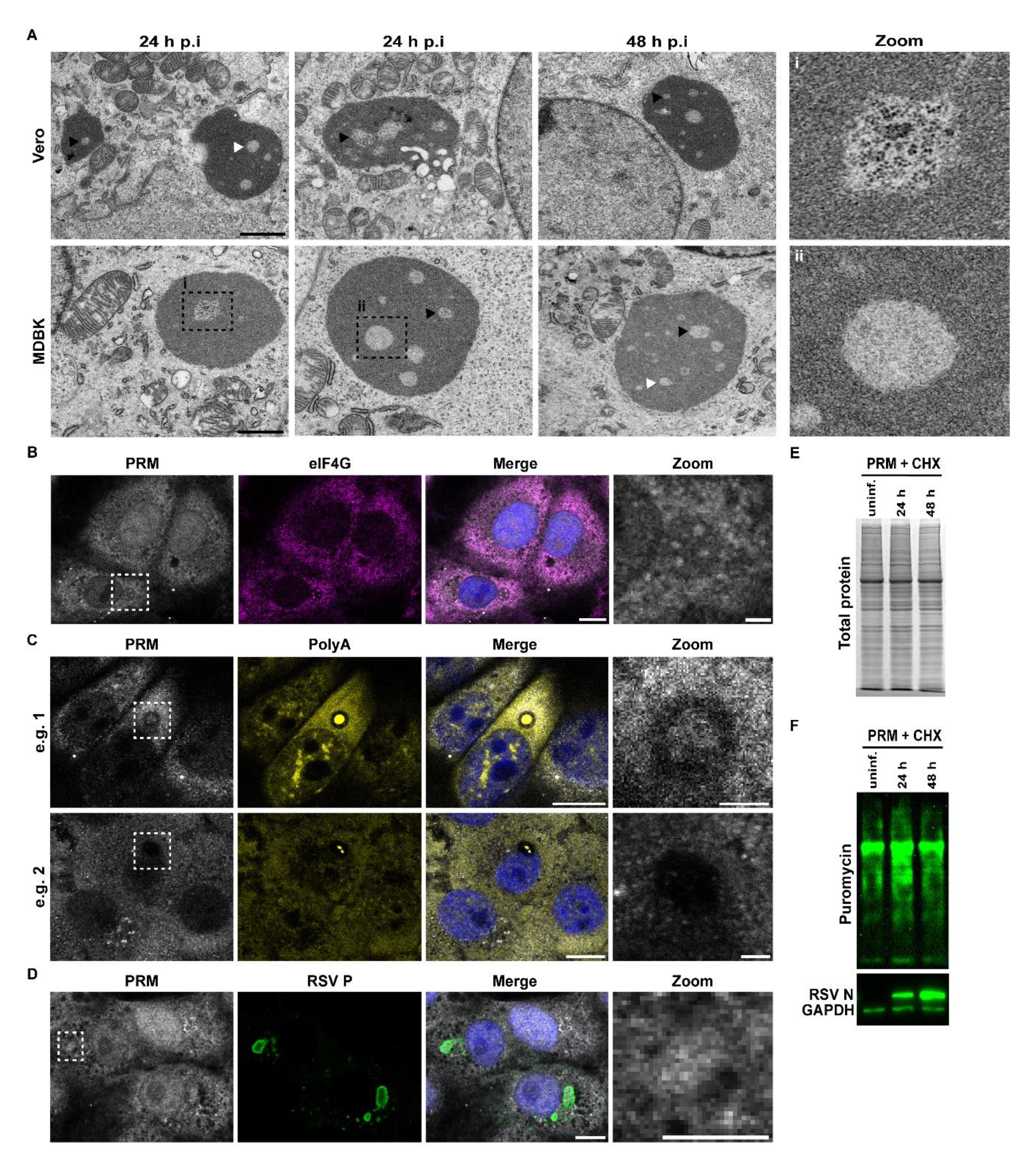
IB microdomains are functional sites containing ribosomes and ribopuromycylated products. (A) Vero and MDBK cells were infected with bRSV for 24 and 48 h, fixed in glutaraldehyde and processed for TEM as detailed in the methods. Representative images are shown from the indicated time points. Scale bars, 1 µm. Black arrow heads indicate IBAGs with concentration of ribosomes and white arrow heads to IBAGs with minimal ribosomal content. Zoom panels are higher magnification images of representative IBAGs boxed in the MDBK panel. (B, C and D) Vero cells infected with bRSV for 24 h were pulsed with puromycin (PRM) for 30 secs to allow incorporation into nascent polypeptides of elongating ribosomes. This is followed by addition of the translation elongation inhibitor cycloheximide (CHX) and incubated for 15 mins at 37°C. Cells were then fixed in PFA and ribosomes with incorporated PRM immuno-stained with an anti-PRM mAb (grey) and FISH staining performed to detect total polyadenylated (PolyA) mRNA (yellow). Immunostaining was also performed for eIF4G (magenta), RSV P (green). Scale bars, 10 µm in main panel and 2 µm in zoom. (E and F) Vero cells left uninfected or infected with bRSV for 24 or 48 h were treated with PRM for 30 secs followed by CHX for 15 mins. Whole cell lysates were then prepared for Coomassie staining to assess total protein levels (E) or levels of ribopuromycylated products by immunoblotting with anti-PRM mAb. RSV N was detected to confirm infection and GAPDH as internal loading control.

### M2-1 associates with eIF4G and drives it’s sub-IB localisation, but only in the context of infection

Finally, we sought to decipher the mechanism of eIF4F recruitment and spatial organisation to these distinct IBAG microdomains inside RSV IBs. We observed that in addition to concentration in IBAGs, eIF4G sometimes also accumulated at the IB periphery, where RSV P (Fig. 6A), N (Fig. 2A) and M2-1 (Fig.4A) can also be found, suggesting an interaction with one or more of these viral proteins. All three viral proteins have been shown to be involved in the recruitment of clients to RSV IBs, e.g. N protein-MDA5 [25], M2-1-PABPC1 [32] and P-RSV M2-1/PP1 [22] interactions. However, the significant co-localisation of M2-1 and eIF4F proteins in IBAGs suggested an M2-1 mediated mechanism by which this translation initiation complex is recruited into IBAGs. We have previously identified N and P co-expression as the minimal components for LLPS, with the resultant ‘pseudo-IBs’ recapitulating the p65 recruitment observed during infection [18]. Here, we used a similar system to study eIF4G recruitment. We initially found that unlike in infected cells, eIF4G remained at the periphery of pseudo IBs formed following the co-expression of N and P alone (Fig. 6B). This was consistent and correlated with an absence of IBAGs and the exclusion of polyA signals from pseudo IBs, presumably since there is no active viral replication in this system (Fig. 6C). However, addition of ectopically expressed M2-1 led to the concentration and co-localisation of both M2-1 and eIF4G at the IB periphery, albeit without subsequent IBAG formation (Fig. 6D). Of note, two populations of condensates were observed following the coexpression of RSV N, P and M2-1. Larger pseudo-IBs that contained eIF4G, M2-1 and P and a smaller population that were devoid of P (indicated by white arrows in images and black arrow in left graph; Fig. 6D). These were found in close proximity to the IBs and later identified as stress granules (SGs), using SG markers G3BP1 (supplementary Fig. 3E) and eIF3A (supplementary Fig. 3F). Together, these findings suggest that spatial reorganisation of the IB into biphasic functional sites requires components only present during infection; however, M2-1 can sequester eIF4G to the peripheral ring of phase separated pseudo-inclusions

**Figure 6.**
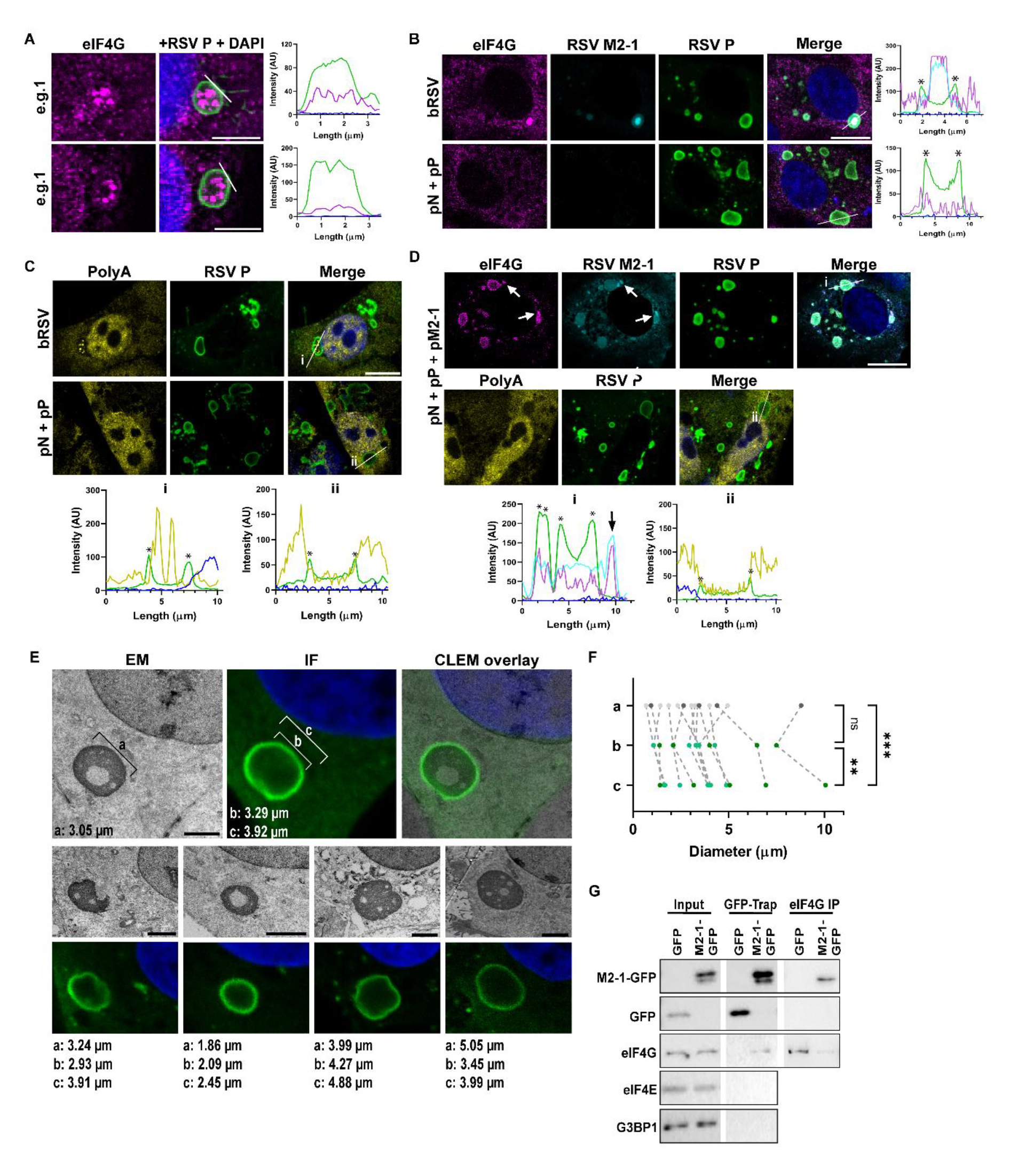
The M2-1 protein mediates eIF4G recruitment into IBAGs. (A) Representative images of bRSV infected Vero cells immuno-stained for eIF4G (magenta) and RSV P (green). Scale bars, 5 µm. Graphs show fluorescent intensity profiles along the white lines drawn in the “RSV P + DAPI” panel. (B and C) Vero cells infected with bRSV or transfected with plasmids expressing RSV N and P proteins were fixed and immuno-stained for eIF4G, RSV M2-1 and P, in B or FISH labelled for total PolyA mRNA and immuno-stained for RSV P, in C. (D) Vero cells transfected with plasmids expressing RSV N, P and M2-1 proteins were immuno-stained for eIF4G, RSV M2-1 and P or FISH labelled for total PolyA mRNA and immuno-stained for RSV P. Scale bars, 10 µm. Graphs show fluorescent intensity profiles along the white lines drawn in the “Merge” panels. Asterisks indicate areas of increased P intensity at the edge of the IBs. (E and F) Using CLEM images of MDBK cells infected with bRSV for 24 h, longest diameter of IBs were measured in EM micrographs (a) and compared to measurements of the internal (b) and external (c) diameter of the same IBs imaged by confocal microscopy. Panels show examples of five IBs measured. Scale bars, 2 µm. (F) All IBs measured from this experiment are presented. One-way ANOVA with Tukey’s multiple comparison was used for statistical analysis; \*\*\**P*<0.001, \*\**P*<0.01, ns, non-significant. (G) eIF4G and M2-1 association. 293T cells were transfected with plasmids expressing M2-1-GFP or GFP as control. Cell lysates were prepared at 24 h post transfection and incubated with GFP-Trap resin or Dynabeads pre-bound with anti-eIF4G mAbs. Bound proteins were analysed by SDS-PAGE and immunoblotting for the indicated proteins. Results are representative of immunoblots from two independent experiments for GFP-TRAP pulldowns and three for eIF4G IP.

Parallel live cell analysis of N-GFP and P pseudo-IBs showed they form liquid droplets that undergo fusion (supplementary Fig. 3A) and fission events (supplementary Fig. 3B) consistent with LLPS and liquid organelle formation. Interestingly, detection of N-GFP and P pseudo-IBs using GFP fluorescence shows even distribution of N throughout the IB, whereas immunostaining with anti-N (in the case of both N and P or N-GFP and P pseudo IBs) consistently shows a characteristic ring, as observed during IF of infected cells (sup. Fig. 3C). These N ring observations, made by us and others previously, have been attributed to a lack of N antibody staining inside the IB due to poor epitope accessibility [25]. Since eIF4G and M2-1 appear to be recruited to this area, perhaps prior to their entrance into the IB, we next turned our efforts to characterising the nature of this ring in more detail. When we compared measurements of individual IB diameters from our CLEM analysis, we found that the diameter of the phase separated IBs measured by EM (Fig. 6E; measurement *a*) were significantly more closely related to the internal (Fig. 6E; *b*) than external (Fig. 6E; *c*) diameter of the N ring observed by IF (Fig 6E and F, and supplementary Fig. 3D). These data suggest that the ring of N detected by IF is not actually within the phase separated IB and likely represents a pool of N protein, perhaps N^0^P at the external periphery of the IB. Of note, although the images used in the analysis are from the same cells they were taken separately. Nevertheless, efforts were made to acquire both correlated images at the same z-position (in the median section of the cells), using the size of the nucleus as reference.

Finally, to confirm an interaction between M2-1 and eIF4G, we performed immunoprecipitation (IP) experiments using cell lysates expressing GFP-tagged M2- 1 (M2-1-GFP) or just GFP, as control. In the first approach using GFP-Trap, eIF4G was found in the bound fraction of M2-1-GFP and not in the GFP only control sample (Fig.6G) while eIF4G interactors, eIF4E [29] and G3BP1 [33] were not detected in this fraction. A parallel IP with eIF4G antibody also revealed M2-1-GFP in the bound fraction and not GFP. In summary, these results indicate that an M2-1-eIF4G association underpins the mechanism by which eIF4G and possibly the eIF4F complex are recruited into IBAGs, possibly trafficking through an extra-IB transitionary domain where proteins accumulate before phase separation.

## Discussion

Orthopneumoviruses, like most negative sense RNA viruses, are entirely dependent on the host cell translation machinery. Herein, we have demonstrated that bovine and human RSV modulate this machinery, specifically within IBs that exist as bi-phasic condensates, with differential composition and ultrastructure. Further, we have shown that the viral M2-1 protein plays a pivotal role in the sequestration of cellular components to effect this modulation by trafficking cellular proteins from the cytoplasm and between the phases of the viral liquid organelle (Fig. 6).

Initially, rearrangement of several proteins involved in translation initiation into IBs and IBAGs together with the association of specific ribosomal subunits such as RPL3 indicated compartmentalisation of translation during infection. Later, the presence of ribosomes and active translation within these sites clearly demonstrated frequent compartmentalisation of translation. Thus, it is clear RSV IBs create cellular microenvironments within which mRNA transcription, viral protein synthesis and genome replication are segregated, likely advantaging viral replication while protecting viral factors and replication intermediates from host antiviral defences. Similar compartmentalisation of the translation machinery has been observed during infection with poxviruses [34], African swine fever virus [35] and reoviruses [36], pointing to conserved viral strategies for subversion of the cell. Interestingly, translational activity is frequently globally suppressed as an antiviral response to infection, via mechanisms such as eIF2α phosphorylation and eIF4G cleavage, which impair translation initiation [37]. Indeed, viruses respond by using a diverse range of strategies to selectively subvert cellular translation and/or these antiviral responses towards viral protein synthesis [38-40]. Interestingly, however, during RSV infection we did not observe gross changes to cellular translation kinetics (Fig. 5E), despite our initial assumption that sequestration of translation initiation components into IBAGs would strongly favour translation of viral protein synthesis over cellular proteins. This may reflect the translational output of the immortalised cell line models used in our experiments and we are currently investigating this recruitment in more physiologically relevant airway models. Nevertheless, our data provides important new evidence that spatially connects translation initiation to the previously reported accumulation of nascent viral mRNA transcripts in IBAGs [17, 18] and also helps to shed lights on the biophysical implications of this at the ultrastructural level.

Focusing in on these morphologically distinct micro-domains (that were confirmed by CLEM to be IBAGs [Fig. 4E and 4F]) we propose that these areas of reduced electron density form as a result of their distinct molecular and biophysical composition. Although it is currently unclear whether IBAGs are also phase-separated structures driven by the same physicochemical properties as the larger IBs [41], they largely appear spherical in nature (Fig. 3, 4 and 5A) and were previously shown to be fluid structures [17], two defining features of phase-separated biomolecular condensates [42, 43]. Within the cell it is well established that condensates are frequently formed from networks of RNA and RNA-binding proteins. We propose a distinct composition for the two phases of the biphasic IB: viral mRNA colocalising with M2-1 and eIF4F proteins in IBAGs and viral genomic RNA, N, P, M2-1, and L in the rest of the IB, supported by our data on the functional partitioning of the IB. This compartmentalisation is reminiscent of the spatial organisation of the cellular nucleoli into functionally distinct sites, whereby FCs form sites of transcription [30], consistent with our findings for IBAGs. Furthermore, nucleolar spatial organisation was shown to represent coexistence of multiple liquid phases, separated by differences in the biophysical properties of their constituent macromolecules [44]. The stability of coexisting liquid phases in a ternary complex was shown to be dependent on the presence of a shared component [31]. We suggest RSV M2-1 as the candidate partner shared between the two phases of RSV IBs, however, the exact drivers and significance of RSV IB bi-phasic organisation require detailed further study.

RSV IBs provide a selective environment where viral processes concentrate. Thus, it is not surprising that their composition would be highly regulated. Condensate components may be segregated into “scaffolds”; proteins that initiate nucleation (RSV N and P proteins in the case of IBs [18]) or “clients”; proteins subsequently recruited into the structures [41]. Nucleation proteins tend to contain specific molecular signatures that support the formation of multiple protein-protein and protein-RNA interactions [43, 45]. They may also be involved in the active recruitment of other proteins, for example, M2-1 and PP1 recruitment by the P protein [22]. Our results showed a previously uncharacterised interaction between eIF4G and M2-1, indicative of a mechanism by which eIF4G and its interacting partners [29] (eIF4F components; Fig. 3B, G3BP1; supplementary Fig. 3E, and eIF3A; supplementary Fig. 3F) are recruited from the cytoplasm into IBs, and ultimately IBAGs. Our detailed analysis of CLEM data identified that a ring of N protein (as well as other scaffolds and clients including eIF4G) concentrates on the periphery of the IB, and it is attractive to postulate this is a staging point for proteins about to enter the phase separated organelle. The mechanistic and ultrastructural basis for this LLPS transitionary phase is the subject of further work in our group.

One conflicting aspect of our research remains the determination of the ultimate site for RSV mRNA translation. Whilst we were able to capture and image translation within IBAGs, which we think is associated with ribosomal sequestration in IBAGs, our global analysis of translation still points to a majorly cytoplasmic location for this process, since host cell translation appears unaffected by viral infection (Fig. 5B, C and D). It may be that the sensitivity of our assays prevents the correct assessment of translational frequency in IBAGs and that all of these structures contain some ribosomes, allowing a pioneer round of translation to continue, prior to IBAG release from the IB. Future elucidation of the mechanisms underpinning ribosomal recruitment to IBs and IBAG content release into the cytoplasm may help to improve our understanding of this process.

To conclude, our findings clearly demonstrate compartmentalisation of the initiation phase of cap-dependent translation within RSV IBAGs, micro-domains which have distinct composition and morphology compared to the rest of the IB. We believe the assembly of eIF4F components and ribosomal subunits on nascent viral mRNA in IBAGs could overcome rate limiting steps in viral replication, facilitating the outcompeting of cellular transcripts for available translation machinery, somewhat akin to the cap-snatching mechanism of influenza. This process, which we propose be termed ‘viral p<u>IC</u>-pocketing’ is mechanistically underpinned by the viral M2-1 protein which appears a master-regulator of the cellular translation machinery.

**Supplementary figure 1.**
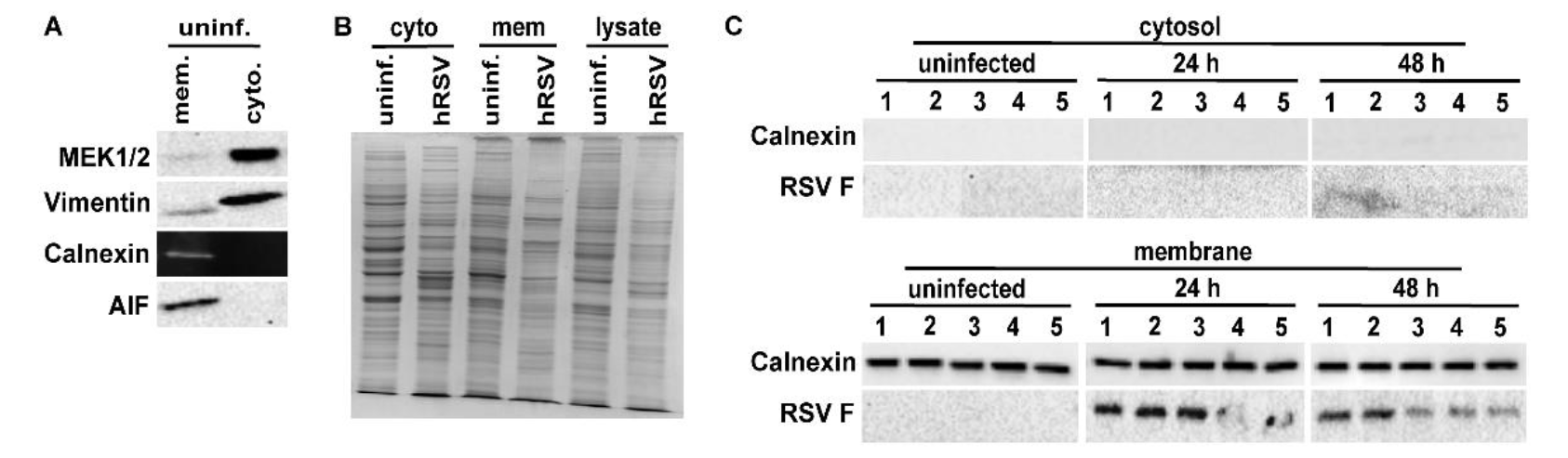
(A) Validation of fractionation kit. Cytosolic and membrane fractions of uninfected A549 cells were separated using a Thermofisher Mem-PER™ Plus membrane protein extraction kit. Immunoblotting was performed on the separated fractions to detect marker proteins of known cellular location. (B) Sub-cellular fractions were made of uninfected or hRSV-infected A549 cells and fractions analysed by SDS-PAGE and coomassie staining. (C) Quintuplicate samples (cytosol and membrane) of fractions analysed by mass spectrometry were immunoblotted for the indicated proteins.

**Supplementary figure 2.**
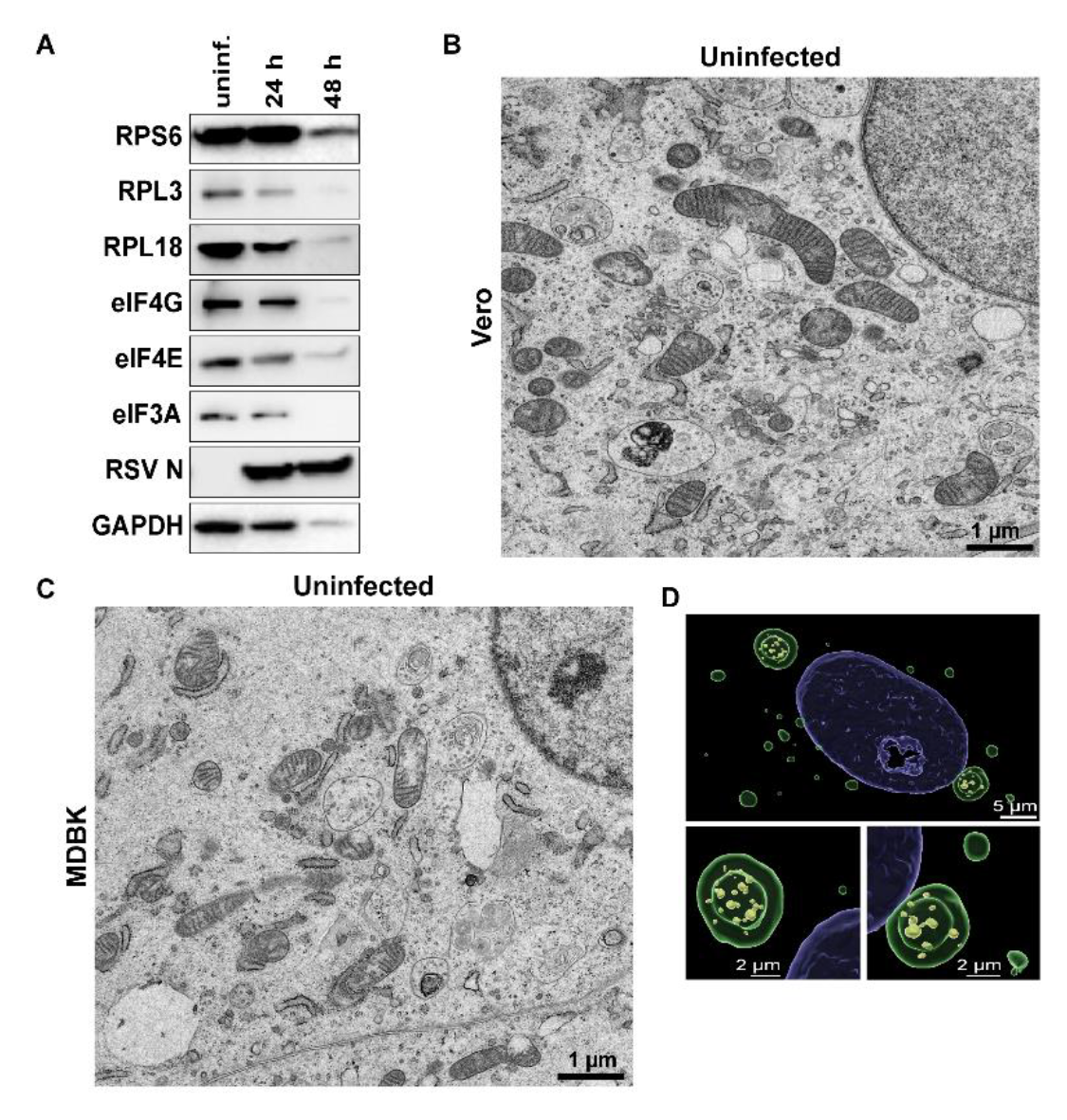
(A) Whole cell lysates prepared from uninfected (uninf.) or from hRSV infected 293T cells at 24 or 48 h p.i, were analysed by SDS-PAGE and immunoblotting for levels of the indicated proteins. (B and C) TEM of uninfected Vero and MDB cells, respectively. (D) Imaris reconstruction of confocal image Z-stacks of bRSV infected Vero cells FISH stained for total PolyA mRNA (yellow) and immuno-stained for RSV P (green). Lower panels are enlarged images of the IBs from the top panel.

**Supplementary figure 3.**
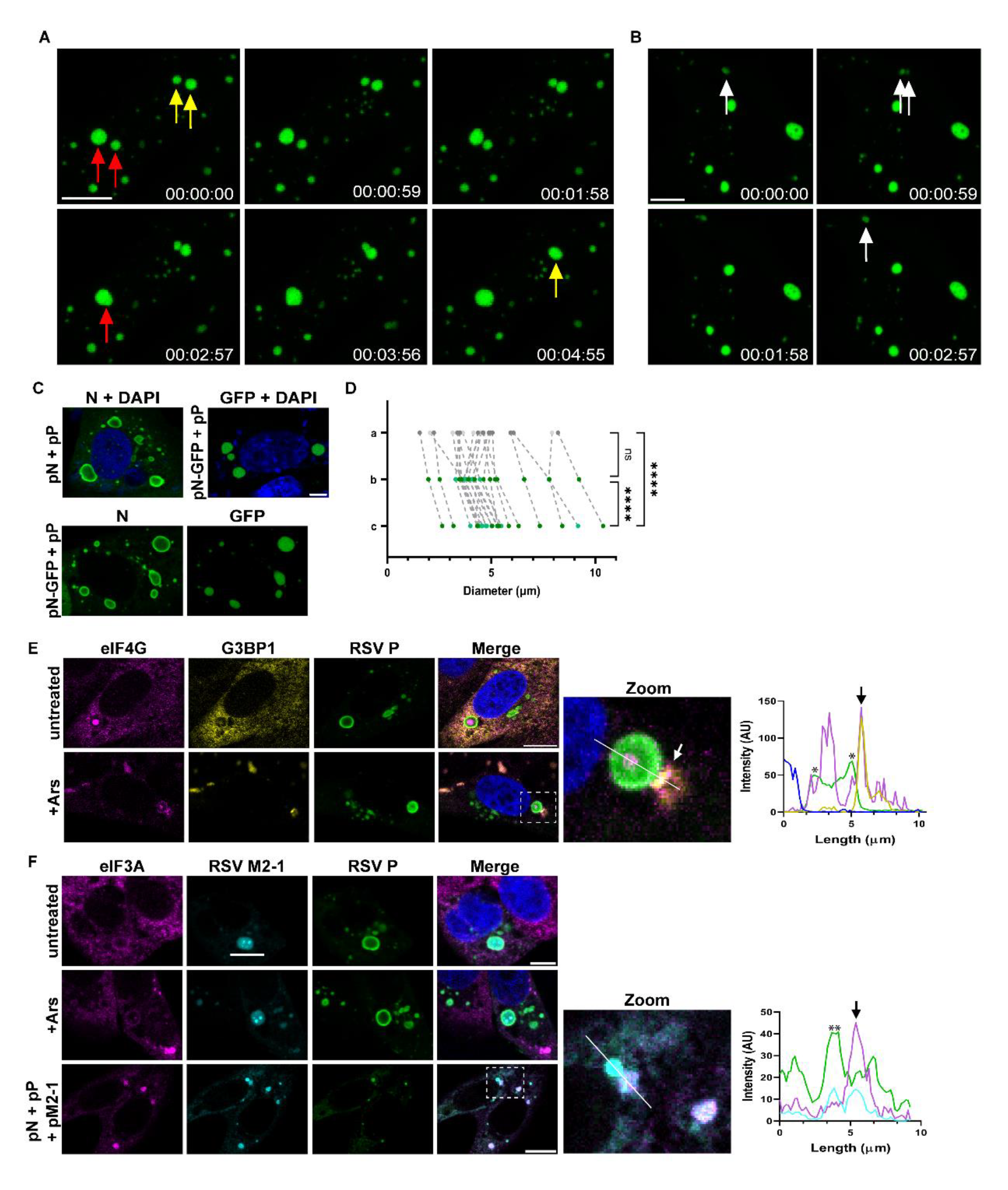
(A and B) IB fusion and fission. Time-lapse imaging of pseudo-IBs (green) formed in Vero cells transfected with plasmids expressing RSV N-GFP and P. Cells were imaged at around 16 h post transfection at 59 sec intervals in a chamber maintained at 37°C, using a Leica Stellaris confocal microscope. Red and yellow arrows in A indicate pseudo-IBs that eventually fuse together. White arrows in B indicates an IB undergoing fission and subsequent fusion. Images are representative of three independent experiments. Scale bars, 10 µm. (C) Pseudo IBs formed in Vero cells transfected with plasmids expressing RSV N and P or N-GFP and P. Detection of RSV N protein with an anti-N mAb shows the characteristic ring pattern whereas, GFP is even distributed through-out the pseudo IB. Scale bar, 5 µm. (D) Comparison of bRSV IB diameters in Vero cells measured from EM and IF images as described in Fig 6E. One-way ANOVA with Tukey’s multiple comparison was used for statistical analysis; \*\*\*\**P*<0.0001, ns; non-significant. (E) bRSV infected Vero cells were treated with 500 µM sodium arsenite (+Ars) for 1 h or left untreated at 24 h p.i. (F) Vero cells were prepared as detailed in E or transfected with plasmids expressing RSV N, P and M2-1 proteins. Cells were then fixed and immuno-stained for eIF4G, G3BP1 and RSV P in E, or for eIF3A, RSV M2-1 and P in F. Scale bars, 10 µm. Enlarged images of the boxed areas are shown in the “zoom” panel. Graphs show fluorescent intensity profiles along the white lines drawn in the “zoom” panel.

## Materials and methods

### Cells and viruses

All cells were cultured at 37°C in a 5% CO2 atmosphere. A549 (human lung epithelial), MDBK (Madin-Darby bovine kidney), Vero (monkey kidney epithelial), 293T (human embryonic kidney) and Hep-2 (human epithelial type 2) cells were obtained from the Pirbright Institute Central Services Unit and maintained in Dulbecco’s modified Eagle’s medium (DMEM; Sigma, Merck) supplemented with 10% heat-inactivated foetal calf serum (FCS; TCS Biologicals), sodium pyruvate (Gibco), and penicillin and streptomycin (Merck). Recombinant bRSV was produced by reverse genetics from bRSV strain A51908 variant Atue51908 (GenBank accession no. <u>AF092942</u>) [46] and hRSV from subtype A (A2 strain). Virus stocks were respectively grown in Vero or Hep-2 cells at 37°C and quantified by TCID50 assay.

### Infections and transfections

For virus infections, virus diluted in serum-free medium was adsorbed to cells at MOI 1 or 2 for 90 mins at 37°C in a 5% CO2 atmosphere. Uninfected cells were incubated with serum-free medium. Following adsorption, the inoculum was removed, and cells incubated in medium containing 2% FCS for the indicated times. Plasmids were transfected into cells using 4 µL TransIT-X2 (Geneflow) for every 2 µg DNA according to the manufacturer’s instructions. The plasmids expressing bRSV N (pN) and P (pP) have been previously described [18]. Plasmids expressing bRSV M2-1 (pM2-1), M2-1-GFP (pM2-1-GFP), N-GFP (pN-GFP) and GFP alone (pGFP) were constructed by inserting the codon-optimised coding sequences into pcDNA3.1 using the same standard cloning strategy. All sequences were confirmed by conventional sanger sequencing.

### Antibodies and reagents

Mouse monoclonal antibodies raised against bRSV N (mAb89), P (mAb12), and M2-1 (mAb90) were previously described [47, 48]. Rabbit monoclonal, anti-RPL3 and anti-RPL18 were purchased from Affinity Biosciences. Rabbit anti-RPS6 (2217), anti-eIF4G (2467), anti-eIF4E (2067), anti-eIF3A (3411), anti-eIF4A (2013), anti-eIF4A1 (2490), anti-eIF4B (3592), anti-eIF4H (3469), and anti-GAPDH (5174) were obtained from Cell Signalling Technology (CST). Mouse anti-G3BP-1 was obtained from BD Biosciences, rat anti-GFP from BioLegend, and mouse anti-puromycin and sodium arsenite were purchased from Sigma, Merck. Secondary horseradish peroxidase-linked antibodies were obtained from CST and Alexa Fluor secondary antibodies from Life Technologies.

### Quantitative mass spectrometry

In-gel digestion was similar to the described protocol [49]. Fractionated samples (normalised to 20 µg) were run approx. 1 cm into a 4-12% NuPage gel (Invitrogen) before staining with Coomassie blue (GelCode Blue Safe Protein Stain, Fisher) for 2 hours, then de-stained with ultrapure water. The entire lane length (1mm wide) was excised and cut into smaller pieces (approx. 1mm^3^) before de-staining with 50% acetonitrile (v/v) in 25mM ammonium bicarbonate. Proteins were reduced for 10 mins at 60°C with 10 mM dithiothreitol (Sigma, Merck) in 50 mM ammonium bicarbonate and then alkylated with 55 mM iodoacetamide (Sigma, Merck) in 50 mM ammonium bicarbonate at room temperature for 30 mins in the dark. Gel pieces were washed for 15 mins in 50 mM ammonium bicarbonate and then dehydrated with 100% acetonitrile. Acetonitrile was removed and the gel plugs rehydrated with 0.01 µg/µL proteomic grade trypsin (Thermo) in 50 mM ammonium bicarbonate. Digestion was performed overnight at 37°C. Peptides were extracted with 50% (v/v) acetonitrile, 0.1% TFA (v/v) and the extracts were reduced to dryness using a centrifugal vacuum concentrator (Eppendorf). Peptides were re-suspended in 3% (v/v) methanol, 0.1% (v/v) TFA for NanoLC MS ESI MS/MS analysis.

LC-MS/MS analysis was similar to that described [50]. Peptides were analysed by on-line nanoflow LC using the Ultimate 3000 nano system (Dionex/Thermo Fisher Scientific). Samples were loaded onto a trap column (Acclaim PepMap 100, 2 cm × 75 μm inner diameter, C18, 3 μm, 100 Å) at 9μl /min with an aqueous solution containing 0.1 % (v/v) TFA and 2% (v/v) acetonitrile. After 3 min, the trap column was set in-line an analytical column (Easy-Spray PepMap® RSLC 50 cm × 75 μm inner diameter, C18, 2 μm, 100 Å) fused to a silica nano-electrospray emitter (Dionex). The column was operated at a constant temperature of 35°C and the LC system coupled to a Q-Exactive mass spectrometer (Thermo Fisher Scientific). Chromatography was performed with a buffer system consisting of 0.1% formic acid (buffer A) and 80% acetonitrile in 0.1% formic acid (buffer B). The peptides were separated by a linear gradient of 3.8 – 50% buffer B over 90 mins at a flow rate of 300 nl/min. The Q-Exactive was operated in data-dependent mode with survey scans acquired at a resolution of 70,000 at m/z 200. Scan range was 300 to 2000m/z. Up to the top 10 most abundant isotope patterns with charge states +2 to +5 from the survey scan were selected with an isolation window of 2.0Th and fragmented by higher energy collisional dissociation with normalized collision energies of 30. The maximum ion injection times for the survey scan and the MS/MS scans were 250 and 50ms, respectively, and the ion target value was set to 1E6 for survey scans and 1E5for the MS/MS scans. MS/MS events were acquired at a resolution of 17,500. Repetitive sequencing of peptides was minimized through dynamic exclusion of the sequenced peptides for 20s.

### Data processing and protein identification

All data were imported, aligned, and peak picked using Progenesis QI for Proteomics (Nonlinear Dynamics,version 4.1). Protein identifications were obtained by searching against Homo sapiens (Uniprot, UP000005640, December 2019) or RSV A2 (UP000181262, December 2019) UniProt databases using PEAKS studio 7 software (Bioinformatics Solutions Inc.) [51]. Searches were conducted using a 1% false discovery rate, which was determined using a decoy database strategy. Peptide and fragment ion tolerances were set to 15ppm and 0.02Da respectively. Two missed tryptic cleavages were permitted. Carbamidomethylation (cysteine) was set as a fixed modification. Oxidation (methionine) was set as a variable modification. Normalized abundance values based on the estimation of the amount of proteins present within samples were imported into MetaboAnalyst 4.0 for univariate and multivariate statistical analysis [52] All data were log transformed and unit scaled (whereby each variable was centred and divided by the standard deviation of the variable to convert to a z-score). Principal Components Analysis (PCA) was used to reduce the high-dimensional dataset and to explore initial data structure. Subsequently, different splits of the data (including comparisons based on time, on case-controls and on cytosol versus membrane proteins) were analysed in order to calculated p-values and fold changes. P-values were calculated by student t-test or - in the case of three-way classifications - by ANOVA. Protein signatures for each split of the data were constructed by including markers which met the criteria of having a FDR-adjusted p-value of < 0.01.

### Functional annotation and pathway analysis

Pathway analysis using the various protein signatures for each comparison was performed using ClueGO (Version 2.5.7), a plug-in application in Cytoscape (Version 3.8.0) [53] searching against the Reactome pathway database. Only pathways with FDR-adjusted p-value < 0.01 and with a minimum of 3 proteins per pathway were considered. Pathway enrichment/depletion analysis was done in a two-sided hypergeometric test and using Bonferroni step down correction. The major pathway identifiers (from the biomarker list) characterising each cluster were highlighted in the generated visualisations.

### Immunoblotting analysis

Cells were lysed in SDS sample buffer (BIO-RAD) supplemented with β-mercaptoethanol (Sigma, Merck) and complete mini-EDTA-free protease inhibitors (Roche). Lysates were then boiled for 10 mins, resolved by SDS-PAGE and transferred to nitrocellulose membranes. Membranes were then blocked and probed for the indicated proteins. All primary antibody incubations were done overnight at 4°C.

### Fluorescent in situ hybridisation

At 24 h p.i, cells were incubated with medium supplemented with 20 μg/ml actinomycin D (Act D; Merck) for 1 h to inhibit cellular transcription. Cells were then fixed with 4% paraformaldehyde (PFA; Merck) in PBS for 15 min, permeabilized with 0.2% Triton X-100 in PBS for 5 min, before blocking with 1% bovine serum albumin (BSA) (Merck) in PBS supplemented with 4 µg/mL free streptavidin, for 1 h. Cells were then re-fixed with 4% PFA for 10 min and incubated at 37°C overnight in hybridisation mix containing 2x SSC, 10% dextran, 1 mg/mL herring sperm DNA, 20% formamide (all purchased from Sigma, Merck), and 1 µM oligo dT probes, for the detection of total polyadenylated (PolyA) RNA (mRNA). The detection of viral NS1 and N mRNA was done using 10 µM total of a mix of probes (sequences in Table1 1) in the hybridisation mix as above, except with a formamide concentration of 50%. Cells were then washed at 42°C with 2x SSC plus 20% formamide for oligo dT probes and 50% formamide for all other probes. This was followed by washes in 2x SSC, then 1x SSC and finally PBS. All probes were single-stranded DNA oligonucleotides synthesised with biotin at the 3’ end (Integrated DNA Technologies) and were detected using streptavidin-Alexa Fluor 488 conjugate (Life Technologies). Cells were then immuno-stained for viral proteins following the immunofluorescence staining protocol, before imaging by confocal microscopy.

**Table 1.**
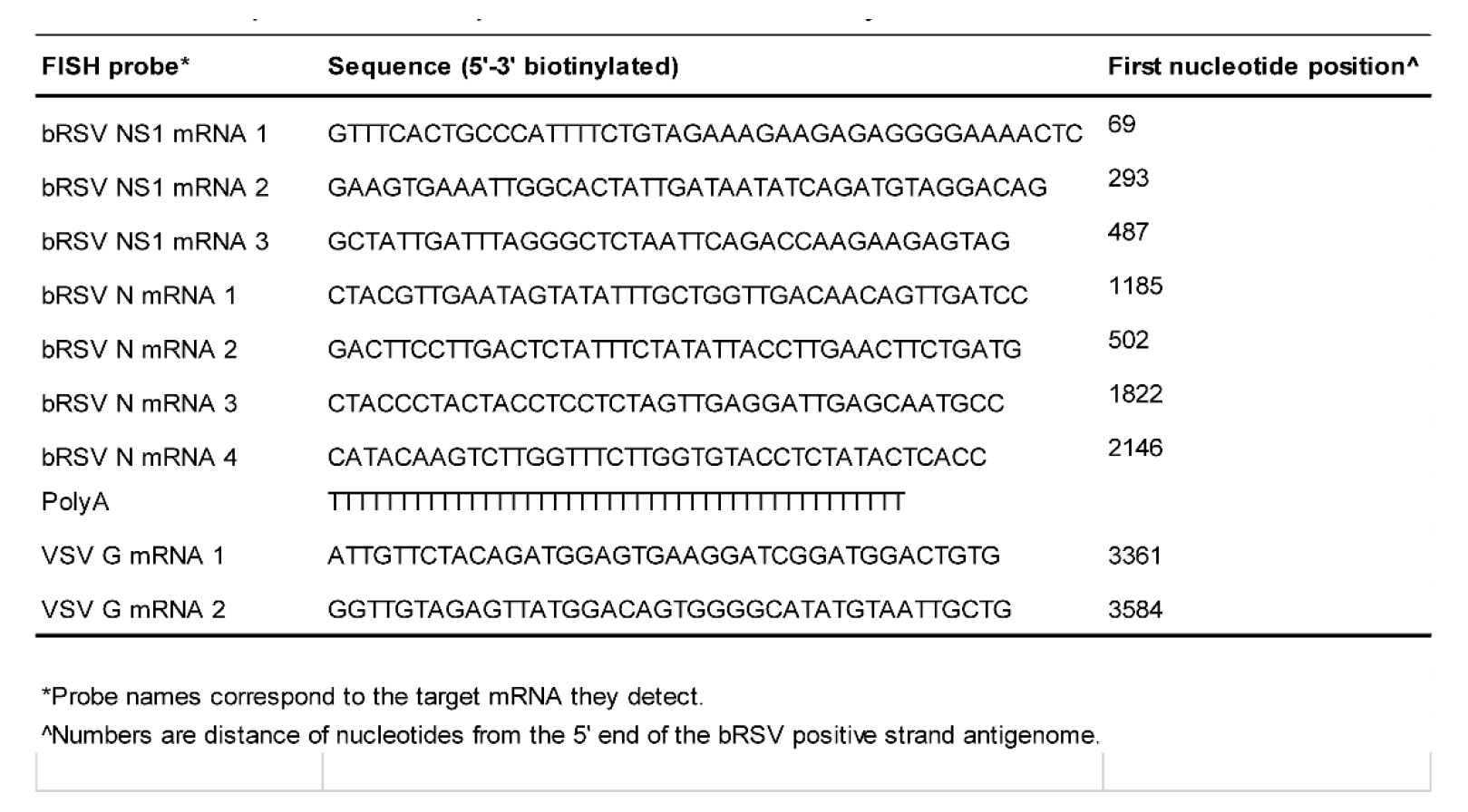
Sequences of the probes used unb FISH analysis

### Confocal immunofluorescence microscopy

Cells were fixed and permeabilised as above before blocking with 1% BSA in PBS for 1 h. Primary antibody incubations were done overnight at 4°C followed by PBS wash and Alexa Fluor secondary antibody (Life Technologies) incubations, for 1 h at room temperature. Cells were then washed and mounted with Fluoromount mounting medium (Invitrogen, ThermoFisher Scientific) containing DAPI for nuclei staining. Fluorescence was imaged on a Leica Stellaris confocal microscope using 405-nm, 488-nm, and 568-nm laser lines for the appropriate dyes and a 63× oil immersion objective. Quantitation of fluorescence signal intensities along line regions of interest were exported from the LAS X software and graphs prepared in GraphPad Prism 9.

### Transmission electron microscopy

Infected cells growing on Thermanox coverslips (ThermoFisher Scientific, UK) were infected and fixed at the indicated times in phosphate-buffered 2% glutaraldehyde (Agar Scientific) for 1 hour followed by 1 hour in aqueous 1% osmium tetroxide (Agar Scientific). Cells were then dehydrated in an ethanol series: 70% for 30 min, 90% for 15 min, and 100% three times for 10 min. A transitional step of 10 min in propylene oxide (Agar Scientific) was undertaken before infiltration with a 50:50 mix of propylene oxide and epoxy resin (Agar Scientific) for 1 hour. After a final infiltration of 100% epoxy resin for 1 hour, the samples were embedded and polymerized overnight at 60°C. Next, 80-μm-thin sections were cut, collected onto copper grids (Agar Scientific), and grid stained using Leica EM AC20 before being imaged at 100 kV in a FEI Tecnai 12 TEM with a TVIPS F214 digital camera.

### Correlative light electron microscopy

Infected cells growing on gridded glass coverslips (MatTek) were fixed and labelled according to the described immunofluorescence method. Selected grid squares were imaged on a Leica TCS SP8 confocal microscope using 405-nm, 488-nm, and 568-nm laser lines for the appropriate dyes. The cells were then fixed in phosphate-buffered 2% glutaraldehyde (Agar Scientific) for 1 hour, 1% osmium tetroxide (Agar Scientific) for 1 hour and 3% uranyl acetate (Agar Scientific) for 15 min before dehydrating in an ethanol series as described above. Next, cells were infiltrated with 100% epoxy resin (Agar Scientific) for 2 hours, embedded and polymerized overnight at 60°C. 80-μm-thin sections were cut from appropriate grid squares, collected unto copper grids (Agar Scientific), and grid stained using a Leica EM AC20 instrument. The specific cells imaged in the confocal were identified and imaged at 100 kV in a FEI Tecnai 12 TEM with a TVIPS F214 digital camera and a ThermoScientific Talos L120C G2 TEM with a Ceta 4K CMOS camera.

### Live imaging

Vero cells were seeded unto Chambered number 1.5 Borosilicate Coverglass slide (Sigma, Merck) before transfection. Cells were transfected with plasmids expressing bRSV N-GFP and P in DMEM supplemented with 2% FCS.16 h post transfection medium was replaced with Leibovitz’s L-15 medium without phenol red (Gibco, ThermoFisher Scientific), and imaged live with a Leica TCS SP8 confocal microscope using a 63× oil immersion objective. During imaging, cells were maintained at 37°C. Z-stacks were imaged for each time series and stacks acquired at intervals of 59 secs.

### Image analysis and statistics

3D reconstruction was performed using Bitplane Imaris software v9.9.1 (Andor Technology PLC, Belfast, UK). The Surface creation tool was used to quantify the number of IBAGs and their volume, using the polyA signal as the source channel. Where required, touching objects were manually split and the ‘magic wand’ option was used to add in objects that had not been properly segmented. The Surface tool was also used to quantify IB volume and longest diameter, using either P or M2-1 signal as the source channel. As there is no direct Surface statistic for diameter within Imaris, the object-oriented (OO) bounding box statistic ‘BoundingBoxOO Length C’ was used to give the length of the longest principal axis, thus the longest diameter of the object. Statistics were exported from Imaris and further processed in GraphPad Prism 9.

### Ribopuromycylation method

This method was performed according to the protocol described [54] with modifications. Cells infected with virus for 24 h, were left untreated or treated with 18.4 μM puromycin (PRM; Sigma, Merck) for 30 secs, followed by incubation at 37°C with 100 μg/mL cycloheximide (CHX; Sigma, Merck) and 18.4 μM PRM for 15 mins. Cells were then fixed, permeabilised and immunostained as described above or cell lysates prepared using Laemmli sample buffer for SDS-PAGE analysis.

### Immunoprecipitation

Cells transiently expressing M2-1-GFP or GFP as control were lysed with cell lysis buffer (CST) supplemented with complete mini-EDTA-free protease inhibitors (Roche) on ice and cell debris removed by centrifugation. Lysates were then incubated at 4°C overnight with GFP-Trap agarose (Chromotek) or Dynabeads (Invitrogen, ThermoFisher Scientific) that were pre-incubated for 30 min at room temperature with anti-eIF4G antibody, following the manufacturer’s protocols. After being washed, samples were eluted in Laemmli buffer supplemented with β-mercaptoethanol, boiled for 10 mins, and resolved by SDS-PAGE.

## Acknowledgements.

This work was supported by a UK Research and Innovation (UKRI; www.ukri.org) Medical Research Council (MRC) New Investigator Research Grant to D.B. (MR/P021735/1) as well as a UKRI Biotechnology and Biological Sciences Research Council (BBSRC; www.ukri.org) Institute Strategic Program Grant (ISPG) to The Pirbright Institute and D.B. (BBS/E/I/00007034 and BBS/E/I/00007030). The funders had no role in study design, data collection and analysis, decision to publish, or preparation of the manuscript.

We acknowledge the support of Trevor Sweeney and Geraldine Taylor (The Pirbright Institute), Nicolas Locker and Matt Brownsword (University of Surrey) and Alex Borodavka (University of Cambridge) for the provision of valuable reagents and technical advice.

